# Effects of probiotics on loperamide-induced constipation in rats

**DOI:** 10.1101/2020.11.23.393843

**Authors:** Takio Inatomi

**Affiliations:** Inatomi Animal Hospital, 1-1-24, Denenchofu, Ota-ku, Tokyo 145-0071, Japan

## Abstract

Constipation, a common health problem, causes discomfort and affects quality of life. This study aimed to evaluate the potential effect of probiotics on loperamide (LP)-induced constipation in Sprague-Dawley (SD) rats, a well-established animal model of spastic constipation. In total, 100 male SD rats (age, 8 weeks; Japan SLC, Inc., Hamamatsu, Japan) were used in the experiments following a 12-day acclimatisation period. They were randomly divided into two treatment groups (groups 1 and 2) of 50 rats each. Spastic constipation was induced via oral administration of LP (3 mg/kg) for 6 days 1 hour before the administration of each test compound. Similarly, a probiotics solution (4 ml/kg body weight) was orally administered to the rats once a day for 6 days in group 2. In group 1, phosphate buffer solution was orally administered once a day for 6 days 1 hour after each LP administration. The changes in body weight, faecal parameters, short-chain fatty acid concentration in faeces, and faecal immunoglobulin (Ig)-A concentration were recorded. In the present study, the oral administration of probiotics improved faecal parameters, short-chain fatty acid concentration in faeces, and faecal IgA concentration. Our results indicate that probiotics increase the levels of intestinal short-chain fatty acids, especially butyric acid, thereby improving constipation and intestinal immunity.

## Introduction

Constipation is a common problem, and probiotics have been reported to improve bowel motility [1,2]. Symptoms of constipation include the following: a decrease in the frequency of bowel movements, decrease in the amount of faeces, painful bowel movements, dry faeces, and dissatisfaction after a bowel movement [3]. Constipation has a significant impact on quality of life, and prebiotics and probiotics are expected to help in the treatment of constipation [4,5]. Therefore, there is a growing interest in better understanding the effects of probiotics on constipation.

Probiotics are defined as living microorganisms that enter the gastrointestinal tract in their active form in sufficient numbers to exert positive effects [6,7]. Probiotics may, in fact, facilitate a return to normal status after a perturbation of the microbiota (e.g. because of the use of antibiotics or illness) or may reduce the degree of change invoked by such challenges. A few studies have measured a probiotic-enhanced return to baseline levels after antibiotic use in humans [8]. The concept of stabilising intestinal microorganisms via the associated probiotics for health improvement dates to the beginning of the last century. Many studies have been conducted to determine the effect of probiotic cultures on health [9–14]. The ingestion of probiotics is beneficial for treating different diarrhoea-like disorders and lowering the levels of metabolites that are harmful to health, including cancerous markers in the colon [15,16]. Probiotic microorganisms promote various immunomodulatory effects by modulating the gut microbiota [17,18]. Thus, probiotic bacteria could be used for the treatment of constipation because of their health-promoting benefits.

Recently, advances in medical treatments have led to increased numbers of immunocompromised patients; some patients contract opportunistic infections caused by some bacterial species, which are considered non-pathogenic bacteria [19]. However, there are only few studies on intestinal immunity during constipation [20]. Therefore, determination of the safety of probiotics is crucial.

Bio-three H (Takeda Consumer Healthcare Company Limited, Tokyo, Japan) has been approved for medical use and is widely used in Japan, China, and India for the treatment and prevention of infectious diseases. In terms of safety, Bio-three H is considered suitable for the treatment of constipation [21].

There have been many studies regarding the role of probiotics in the mitigation of constipation. However, to the best of our knowledge, only few studies have investigated the role of safety-guaranteed probiotics in the mitigation of constipation and gut immunity. Therefore, this study aimed to investigate the effects of safety-guaranteed probiotics on constipation relief and gut immunity in a rat model.

## Materials and methods

### Ethical approval

This study was conducted at Inatomi Animal Clinic in Tokyo Prefecture, Japan. It was performed under the fundamental guidelines for the proper conduct of animal experiments and related activities at academic research institutions under the jurisdiction of the Ministry of Education, Culture, Sports, Science, and Technology. It was approved by the Ethics Committee of the Inatomi Animal Clinic (Tokyo, Japan; approval number 2020-003).

### Experimental animals

One hundred male Sprague-Dawley rats (age, 8 weeks; Japan SLC, Inc., Hamamatsu, Japan) were used in the experiments following a 12-day acclimatisation period. They were randomly divided into two treatment groups (groups 1 and 2) of 50 rats each. Animals were housed individually in a polycarbonate cage, in a temperature (20-23°C)- and humidity (40-50%)-controlled room. The light/dark cycle was 12/12 hours, and basic diet (Rodent Diet CL-2, CLEA JAPAN, Tokyo, Japan) and water were supplied *ad libitum*. The probiotics solution used in this study was Bio-three H. Bio-three H (1 g) contains 50 mg of *Bacillus mesentericus* TO-A, 10 mg of *Enterococcus faecalis* T-110, and 50 mg of *Clostridium butyricum* TO-A. The probiotics solution was dissolved in phosphate buffer solution (PBS) to a final concentration of 12.5 mg/ml. In group 2, the probiotics solution (4 ml/kg body weight) was orally administered once a day for 6 days 1 hour after each loperamide (LP) administration, as suggested by previously reported studies [22,23]. In group 1, a PBS was orally administered once a day for 6 days 1 hour after each LP administration.

### Induction of constipation in the rats

Constipation was induced in all rats through the oral administration of 3 mg/kg of LP, once a day for 6 continuous days at 1 hour before administration of each test material [22–25].

### Changes in body weight

The body weights of individual rats were daily measured starting from 1 day before administration of the test compounds through the sixth day of administration of the test compounds and LP.

### Measurement of faecal parameters

The excreted faecal pellets of individual rats during a 24-hour period were collected 1 day before the first administration of the test compound and immediately after the fourth administration for a duration of 24 hours. The total number, water content, and wet weight of the faecal pellets were measured. The collected faecal pellets were dried at 60°C in a general dry oven for 24 hours to obtain faecal dry weights.

### Short-chain fatty acid concentration in faeces

The short-chain fatty acid (SCFA) concentration in faeces of each rat was measured using gas chromatography, as described previously [26]. Approximately 0.5 g of faeces from the dissections described above was gently squeezed into a micro-centrifuge tube containing 1 mL of 10% meta-phosphoric acid with 0.4 mL of 4-methyl valeric acid per mL added as an internal standard. The solution was thoroughly mixed using a vortex mixer and centrifuged at 5,700 ×g for 20 minutes at 4°C. The SCFA content of the supernatant was measured using an HP Agilent 6890 series gas chromatograph (Agilent Technologies Inc., Santa Clara, CA, USA) fitted with an HP 5973 series mass spectrometer (Agilent Technologies Inc.). The columns (Agilent Technologies) used were HP-free fatty acid polyester stationary phase capillary columns of polyethylene glycol on Shimalite TPA 60/80, measuring 30-m long with a 0.25-mm internal diameter.

The parameters of gas chromatography were as follows: 1-µl injection volume, 240°C injector temperature, 12.15 psi pressure, and 1.1 mL min-1 constant flow using helium as a carrier. Fatty acids were eluted with the following oven program: 80°C initial temperature hold for 5 minutes, ramp 10°C min−1 to 240°C, and hold for 12 minutes. Individual SCFA concentrations were expressed in mg/g wet faeces.

### Faecal immunoglobulin A concentration

Faeces were suspended in six times the weight of PBS and extracted at 25°C for 24 hours. The extract was centrifuged at 1500 ×g for 10 minutes to collect the supernatant and stored at −30°C, and then, immunoglobulin (Ig) A was measured using an enzyme-linked immunosorbent assay (ELISA) quantitation kit (Betyl Laboratories, Montgomery, Texas, USA). ELISA was conducted according to the manufacturer protocol.

Briefly, 100 μl of the sample or standard was added to the wells of the plate and allowed to stand at 22°C for 1 hour; the plate was then washed 4 times. Next, a chicken IgA detection antibody was added to each well and allowed to stand at 22°C for 1 hour; it was then washed 4 times. Horseradish peroxidase solution A was allowed to stand at 22°C for 30 minutes, and the plate was then washed 4 times. One hundred microliters of 3,3',5, 5'-tetramethylbenzidine substrate solution was added to each well, and the plate was developed in the dark for 30 minutes at 22°C. After stopping the reaction by adding 100 μl of the stop solution to each well, the absorbance was measured at 45-nm wavelength using a plate reader. Subsequently, a calibration curve was prepared, and the IgA concentration was calculated.

### Statistical analysis

The Kolmogorov-Smirnov test was used to assess the normal distribution of the data before statistical analysis was performed. Mann-Whitney U tests were performed using EZR software (Saitama Medical Center, Jichi Medical University); EZR is a graphical user interface for R (The R Foundation for Statistical Computing, version 2.13.0). The significance level was set at *p*<0.05.

## Results

### Effect on body weight

Changes in body weight are shown in Table 1. There were no differences in body weight between group 1 and group 2 during this study.

**Table 1.** Changes in body weight of the rats (g).

|  | Day -1 | Day 1 | Day 6 |
| --- | --- | --- | --- |
| Group 1 | 290.1±1.641 | 287.3±1.776 | 290.9±2.236 |
| Group 2 | 292.5±1.552 | 291.2±1.803 | 288.5±1.941 |

### Changes in faecal parameters

Faecal parameters are shown in Tables 2 and 3.

**Table 2.** Faecal parameters on day −1.

|  | Pellet numbers<br>(numbers/rat) | Wet weights of<br>faeces (g/rat) | Dry weights of<br>faeces (g/rat) | Water contents of<br>faeces (%/rat) |
| --- | --- | --- | --- | --- |
| Group 1 | 58.0±0.42 | 9.41±0.15 | 7.54±0.16 | 29.9±0.27 |
| Group 2 | 57.7±0.45 | 9.34±0.15 | 7.41±0.2 | 30.1±0.24 |

**Table 3.** Faecal parameters on day 4.

|  | Pellet numbers<br>(numbers/rat) | Wet weights of<br>faeces (g/rat) | Dry weights of<br>faeces (g/rat) | Water contents of<br>faeces (%/rat) |
| --- | --- | --- | --- | --- |
| Group 1 | 30.0±0.19<sup>a</sup> | 4.12±0.05<sup>a</sup> | 3.55±0.04<sup>a</sup> | 13.9±0.39<sup>a</sup> |
| Group 2 | 58.3±0.29<sup>b</sup> | 9.01±0.10<sup>b</sup> | 6.70±0.06<sup>b</sup> | 25.4±0.75<sup>b</sup> |

<sup>a, b</sup> Different letters within columns indicate differences between the treatment groups ( $p<0.01$ ).

At 1 day before treatment for 24 hours, there were no differences in pellet numbers, wet weights of faeces, dry weights of faeces, and water contents of faeces between group 1 and group 2. After the fourth administration for 24 hours, pellet numbers, wet weights of faeces, dry weights of faeces, and water contents of faeces were significantly higher in group 2 than in group 1 (*p*<0.01).

### Short-chain fatty acid concentration in faeces

Short-chain fatty acid concentrations in faeces are shown in Table 4.

**Table 4.** Short-chain fatty acid concentration in faeces (mg/g).

|  | Day -1 |  | Day 4 |  |
| --- | --- | --- | --- | --- |
|  | n-butyric acid<br>(mg/g) | acetic acid<br>(mg/g) | n-butyric acid<br>(mg/g) | acetic acid<br>(mg/g) |
| Group 1 | 0.497±0.007 | 1.481±0.016 | 0.448±0.007<sup>a</sup> | 1.480±0.015<sup>a</sup> |
| Group 2 | 0.489±0.007 | 1.519±0.015 | 0.523±0.007<sup>b</sup> | 1.561±0.020<sup>b</sup> |

<sup>a, b</sup> Different letters within the columns indicate differences between the treatment groups ( $p<0.01$ ).

At 1 day before treatment for 24 hours, there were no differences in n-butyric and acetic acid concentrations in faeces between group 1 and group 2. After the fourth administration for 24 hours, n-butyric and acetic acid concentrations in faeces were significantly higher in group 2 than in group 1 (*p*<0.01).

### Faecal IgA concentration

Faecal IgA concentrations are shown in Table 5.

**Table 5.** Faecal immunoglobulin A concentration (mg/g).

|  | Day -1 | Day 4 |
| --- | --- | --- |
| Group 1 | 2.05±0.04 | 2.04±0.04<sup>a</sup> |
| Group 2 | 2.04±0.04 | 2.36±0.03<sup>b</sup> |

<sup>a, b</sup> Different letters within the columns indicate differences between the treatment groups ( $p<0.01$ ).

At 1 day before treatment for 24 hours, there were no differences in IgA concentrations in faeces between group 1 and group 2. After the fourth administration for 24 hours, IgA concentrations in faeces were significantly higher in group 2 than in group 1 (*p*<0.01).

## Discussion

In the present study, the oral administration of Bio-three H improved faecal parameters, short-chain fatty acid concentration in faeces, and faecal IgA concentration. Constipation can arise owing to various causes, including dietary habits; use of chemical compounds, such as morphine; and psychological stress [27]. In this study, the dose of Bio-three H used in rats was determined based on the dose used in humans. The administration of Bio-three H did not show any adverse effects on rats. The results of the faecal parameters of group 1 24 hours after the fourth administration suggested that constipation was properly induced using LP, in accordance with previous studies [22–25]. The body weight of rats was not markedly different between groups 1 and 2 in this study. These results are consistent with those of previous studies [22,23].

Shi et al. reported significantly reduced faecal levels of SCFA (acetic acid, propionic acid, and butyric acid) in a population with constipation compared with those in a healthy population [28]. These observations indicate an association between the occurrence of constipation and the intestinal levels of SCFAs. Therefore, SCFAs produced by intestinal flora or probiotics were believed to be effective for constipation alleviation. Moreover, it was reported that the administration of *Lactobacillus plantarum* NCU116 significantly improved symptoms of constipation in mice and led to significant increases in acetic acid and propionic acid levels in their faeces [29].

SCFA stimulates the intestinal tract and enhances motility as one of the mechanisms of action of SCFA on constipation [30,31]. In this study, Bio-three H significantly increased the faecal concentrations of butyric acid and acetic acid in LP-induced constipation model rats. These results are consistent with those of our previous study [32]. In support of past studies, the current study findings suggest that three bacteria contained in Bio-three H produced SCFA. Bio-three H significantly improved faecal parameters, such as pellet numbers, wet weights of faeces, dry weights of faeces, and water contents of faeces, in this study. From the above results, it is considered that the administration of Bio-three H improved constipation symptoms by increasing SCFA production, and as a result, the faecal parameters were improved.

Havenaar and Spanhaak demonstrated that probiotics stimulate the immunity of animals in two ways: 1) flora from the probiotic migrate throughout the gut wall and multiply to a limited extent; and 2) antigens released by dead microorganisms are absorbed, thus stimulating the immune system [33]. Probiotics containing *B. mesentericus* TO-A, *C. butyricum* TO-A, and *S. faecalis* T-110 cause an increase in immunoglobulin production from the mesenteric lymph nodes in rats [34], stimulate the T-helper 1 immune response in peripheral blood mononuclear and dendritic cells [35], and cause an influx of CD8+ T cells into the intestinal mucosa. These changes may enhance intestinal immunity by CD8+ T cells in young chicks [36] and increase IgA concentrations in the jejunum and ileum in broiler chickens [37]. IgA plays an important role in intestinal immunity by binding to and neutralising pathogens and toxins in the intestinal tract [38]. Intestinal IgA is mainly produced by Peyer plate plasmatic cells, and IgA production by plasmatic cells is induced by follicular helper T cells [39]. The follicular helper T cells differentiate from regulatory T cells in the intestinal tract, and regulatory T cells differentiate from naive T cells [40]. Recently, it has also been shown that butyric acid promotes the differentiation of naive to regulatory T cells in the intestinal tract [41]. In this study, the administration of probiotics increased faecal IgA concentrations. This is because probiotics promoted the differentiation of naive T cells into regulatory T cells by producing butyrate in the intestinal tract, and the increased regulatory T cells differentiated into follicular T cells, which may have indirectly increased IgA production. In support of past studies, the current study findings suggest that three bacteria contained in Bio-three H increased faecal IgA concentrations. However, the mechanism by which this probiotic contributes to the increase in the butyrate concentration in the intestinal tract and stimulates IgA production, as well as the relationship between constipation and IgA production, needs to be studied in detail in the future.

## Conclusions

Our results indicate that probiotics containing *B. mesentericus* TO-A, *E. faecalis* T-110, and *C. butyricum* TO-A increase the levels of intestinal SCFA, especially butyric acid, thereby improving constipation and intestinal immunity. In this study, the probiotics improved constipation symptoms and immune status in rats; however, their effect in humans is unknown. In addition, this study showed the effect of the probiotics on loperamide-induced constipation, but the effect on constipation caused by other factors is unknown. Further studies in humans are needed, but the results of the current study show the potential of this probiotic in improving constipation and immune status in immunosuppressed humans.

## Author Contributions

The authors have performed everything related to this study.

## References

1. Gallagher PF, O’Mahony D, Quigley EM. Management of chronic constipation in the elderly. Drugs Aging. 2008;25: 807–821.

2. Müller-lissner S, Rykx A, Kerstens R, Vandeplassche L. A double-blind, placebo-controlled study of prucalopride in elderly patients with chronic constipation. Neurogastroenterol Motil. 2010;22: 991.

3. Beery RMM, Chokshi RV. Overview of constipation. New York: Springer; 2014.

4. Rothbarth J, Bemelman WA, Meijerink WJ, Stiggelbout AM, Zwinderman AH, Buyze-Westerweel ME, et al. What is the impact of fecal incontinence on quality of life? Dis Colon Rectum. 2010;44: 67–71.

5. Whitehead WE, Wald A, Norton NJ. Treatment options for fecal incontinence. Dis Colon Rectum. 2001;44: 131–142.

6. Heller KJ. Probiotic bacteria in fermented foods: Product characteristics and starter organisms. Am. J Clinic Nutr. 2001;73: 374s–379s.

7. Sanders ME. Probiotics: Definition, sources, selection, and uses. Clin Infect Dis. 2008;46: S58–S61.

8. Engelbrektson AL, Korzenik JR, Sanders ME, Clement BG, Leyer G, Klaenhammer TR, et al. Analysis of treatment effects on the microbial ecology of the human intestine. FEMS Microbiol Ecol. 2006;57:239–250.

9. Bezkorovainy A. Probiotics: determinants of survival and growth in the gut. Am J Clinical Nutr. 2001;73: 399s–405s.

10. Fuller R. History and development of probiotics. New York: Springer; 1992.

11. Isolauri E. Probiotics in human disease. Am J Clinic Nutr. 2001;73: 1142S–1146S.

12. Kalliomäki M, Salminen S, Arvilommi H, Kero P, Koskinen P, Isolauri E. Probiotics in primary prevention of atopic disease: A randomised placebo-controlled trial. Lancet. 2001;357: 1076–1079.

13. Kopp-Hoolihan L. Prophylactic and therapeutic uses of probiotics: a review. J Am Diet Assoc. 2001;101: 229–241.

14. Spanhaak S, Havenaar R, Schaafsma G. The effect of consumption of milk fermented by Lactobacillus casei strain Shirota on the intestinal microflora and immune parameters in humans. Eur J Clin Nutr. 1998;52: 899–907.

15. De Roos NM, Katan MB. Effects of probiotic bacteria on diarrhea, lipid metabolism, and carcinogenesis: A review of papers published between 1988 and 1998. Am J Clinical Nutr. 2000;71: 405–411.

16. De Vrese M, Marteau PR. Probiotics and prebiotics: Effects on diarrhea. J Nutr. 2007;137; 803S–811S.

17. Isolauri E, Kirjavainen P, Salminen S. Probiotics: A role in the treatment of intestinal infection and inflammation? Gut. 2002;50: iii54–iii59.

18. Mazmanian SK, Round JL, Kasper DL. A microbial symbiosis factor prevents intestinal inflammatory disease. Nature. 2008;453: 620.

19. Di Rosa R, Creti R, Venditti M, D’Amelio R, Arciola CR, Montanaro L, et al. Relationship between biofilm formation, the enterococcal surface protein (Esp) and gelatinase in clinical isolates of Enterococcus faecalis and Enterococcus faecium. FEMS Microbiol Lett. 2006;256: 145–150.

20. Khalif IL, Quigley EMM, Konovitch EA, Maximova ID. Alterations in the colonic flora and intestinal permeability and evidence of immune activation in chronic constipation Dig Liver Dis. 2005;37: 838–849. doi: 10.1016/j.dld.2005.06.008.

21. Noguchi N, Nakaminami H, Nakase K, Sasatsu M. Characterization of *Enterococcus* strains contained in probiotic products. Biol Pharm Bull. 2011;34: 1469–1473.

22. Choi JS, Kim JW, Cho HR, Kim KY, Lee JK, Sohn JH, et al. Laxative effects of fermented rice extract in rats with loperamide-induced constipation. Exp Ther Med. 2014;8: 1847–1854.

23. Choi JS, Kim JW, Kim KY, Lee JK, Sohn JH, Ku SK. Synergistic effect of fermented rice extracts on the probiotic and laxative properties of yoghurt in rats with loperamide-induced constipation. Evid Based Complement Alternat Med. 2014;2014: 878503.

24. Bustos D, Ogawa K, Pons S, Soriano E, Banji JC, Bustos FL. Effect of loperamide and bisacodyl on intestinal transit time, fecal weight and short chain fatty acid excretion in the rat. Acta Gastroenterol Latinoam. 1991;21: 3–9.

25. Wintola OA, Sunmonu TO, Afolayan AJ. The effect of Aloe ferox mill. In the treatment of loperamide-induced constipation in Wistar rats. BMC Gastroenterol. 2010;10: 95.

26. Murugesan GR, Romero LF, Persia ME. Effects of protease,phytase and a Bacillus sp. direct-fed microbial on nutrient and energy digestibility, ileal brush border digestive enzyme activity and caecal short-chain fatty acid concentration in broiler chickens. Plos One. 2014;9: p.e101888. doi: 10.1371/journal.pone.0101888.

27. Kakino M, Tazawa S, Maruyama H, Tsuruma K, Araki Y, Shimazawa M, et al. Laxative effects of agarwood on low-fiber diet-induced constipation in rats. BMC Complement Altern Med. 2010;10: 68.

28. Shi Y, Chen Q, Huang Y. Function and clinical implications of short-chain fatty acids in patients with mixed refractory constipation. Colorectal Dis. 2016;18: 803–810. doi: 10.1111/codi.13314.

29. Li C, Nie SP, Zhu KX, Xiong T, Li C, Gong J, et al. Effect of Lactobacillus plantarumNCU116 on loperamide-induced constipation in mice. Int J Food Sci Nutr. 2015;66: 533–538. doi: 10.3109/09637486.2015.1024204.

30. Kamath PS, Phillips SF. Initiation of motility in canine ileum by short chain fatty acids and inhibition by pharmacological agents. Gut. 1988;29: 941–948. doi: 10.1136/gut.29.7.941.

31. Kamath PS, Phillips SF, Zinsmeister AR. Short-chain fatty acids stimulate ileal motility in humans Gastroenterology. 1988;95: 1496–1502. doi: 10.1016/s0016-5085(88)80068-4.

32. Inatomi T. Laying performance, immunity and digestive health of layer chickens fed diets containing a combination of three probiotics. Science Postprint. 2016;1: e00058. doi: 10.14340/spp.2016.03A0001.

33. Havenaar R, Spanhaak S. Probiotics from an immunological point of view. Curr Opin Biotechnol. 1994;5: 320–325. doi:10.1016/0958-1669(94)90036-1.

34. Okabe M, Matsuo A, Nishida E, Tachibana H, Yamada K. Dietary effect of a live-bacterial drug on lipid metabolism and immune function of sprague-dawley rats [article in Japanese]. Nippon Shokuhin Kagaku Kogaku Kaishi (J Jpn Soc Food Sci). 2003;50: 224–229.

35. Hua MC, Lin TY, Lai MW, Kong MS, Chang HJ, Chen CC. Probiotic Bio-Three induces Th1 and anti-inflammatory effects in PBMC and dendritic cells. World J Gastroenterol. 2010;16: 3529–3540.

36. Huang A, Shibata E, Nishimura H, Igarashi Y, Isobe N, Yoshimura Y. Effects of probiotics on the localization of T cell subsets in the intestine of broiler chicks. J Poult Sci. 2013;50: 275–281.

37. Inatomi T. Growth performance, gut mucosal immunity and carcass and intramuscular fat of broilers fed diets containing a combination of three probiotics. Sci Postprint. 2015;1: e00052. doi: 10.14340/spp.2015.10A0001.

38. Xiong N, Hu S. Regulation of intestinal IgA responses. Cell Mol Life Sci. 2015;72: 2645–2655.

39. Winstead CJ. Follicular helper T cell-mediated mucosalbarrier maintenance. Immunol Lett. 2014;162: 39–47.

40. Tsuji M, Komatsu N, Kawamoto S, Suzuki K, Kanagawa O, Honjo T, et al. Preferential generation of follicular B helper T cells from Foxp3+ T cells in gut Peyer’s patches. Science. 2009;323: 1488–1492.

41. Furusawa Y, Obata Y, Fukuda S, Endo TA, Nakato G, Takahashi D, et al. Commensal microbe-derived butyrate induces the differentiation of colonic regulator y T cells. Nature. 2013;504: 446–450.

